# Age-dependent effects of reduced mTor signalling on life expectancy through distinct physiology

**DOI:** 10.1101/719096

**Authors:** Mirre J P Simons, Laura Hartshorne, Suzan Trooster, Jessica Thomson, Marc Tatar

## Abstract

Research on the mechanisms of ageing has identified ways via which lifespan can be extended in model organisms, increasing the potential for translation of these findings to our own species. However, the large majority of research on animal models involves dietary, genetic or pharmacological treatments throughout life – limiting translational potential and ignoring age-dependent effects. Previously, we have suggested using demographic meta-analysis that reduced mTor signalling has the potential to instantly rejuvenate. We have now tested this prediction experimentally using large-scale demographic data (N > 10,000) combined with conditional knockdown of mTor in *Drosophila melanogaster*. Indeed, reduced mTor decreased mortality rate when applied during old age. Interestingly, we found that transient treatment during early adult life had long-lasting benefits. Age-dependent deep-RNAseq indicated that these effects arose from distinct physiology and implicate alternative splicing as a potential mechanism for the long-lasting benefits of transient mTor reduction. These findings suggest that reducing mTor short term or during old age could be used to combat ageing. In addition, our findings suggest that the results from experimental research on mTor signalling, and potentially other mechanisms of ageing, that employ life-long interventions are likely to be a complex composite of age-dependent effects that counteract or enhance each other.

## Introduction

The biology of ageing field has progressed our understanding of mechanisms and interventions that can extend health- and lifespan. The most potent and researched of these are dietary restriction^1–3^ and reducing mTor (mechanistic Target of Rapamycin) signalling^4–6^. These interventions we now (only) partially understand and there is a growing wish and reality for translation to our own species^7^. A key factor that hinders the translation of these findings to humans is however that the large majority of the experiments in animal models are carried out for their entire lifetimes, which begs the question whether long-term treatment will be required in humans as well. Any long-term treatment will be hard to apply in our own species and close to impossible to study in clinical trials. More immediate benefits to health- and lifespan are sought for to hold translational promise^8,9^. Notably, dietary restriction can have instant benefits on health indicators^2,10^, but trials in humans have only been conducted over a relatively short timespan^10,11^. Animal experiments have shown that in terms of longevity, mortality risk is instantly modulated by diet in flies^12–14^, but experiments in other organisms, namely rodents suggest late-life treatment is not necessarily pro-longevity^15,16^. In contrast, short-term energy restriction in early life can result in long-lasting health benefits, as for example in mice by restricting milk access^17^.

Reduced mTor signalling has now also been suggested to improve life expectancy after short-term treatment or in old animals. Rapamycin treatment from 600 days of age onwards resulted in longevity benefits (especially in females, as control biased mortality occurred prior to drug treatment in males)^9^. Recently, short term transient rapamycin treatment (of 3 months) of mice aged 2 years led to a lifespan extension, but the authors acknowledge sample size for this study is limited^18^. We have recently also argued using demographic meta-analysis across four model species that reducing mTor signalling at old age could reduce mortality risk^4^. Such inference from demographic models can be informative^4,15,16,19,20^, but only by testing different ages of manipulation can such demographic patterns be tested for causality^8,12,13,21^. Here, we present the first of such comprehensive evidence using large-scale demography comprising over 10,000 individual flies (*Drosophila melanogaster*), showing that reduction of mTor in late life results in instant benefits on life expectancy. In addition, we find that short transient mTor knockdown during early adulthood has long-lasting benefits reducing mortality in late life even when mTor levels are back to unmanipulated levels. Age-dependent RNAseq data from this experiment showed that these two effects originate from differential physiology, potentially derived from differences in alternative splicing.

## Results and discussion

### Experimental design and GeneSwitch kinetics

Age-dependent knockdown of mTor (using *in vivo* RNAi^22^) was achieved using the well-established conditional GeneSwitch system^23,24^, by feeding adult flies mifepristone (RU486). We chose RNAi over rapamycin treatment as genetic suppression of mTor signalling results in larger effects^4^ and more potent rapalogs are currently being developed^25^. The ligand RU486 for GeneSwitch allows close experimental control of downstream genetic tools. Our experimental design consisted of four groups. A transient treatment with flies fed food containing RU486 for 12 days during early adult life (starting at age 15±1), a late life group fed RU486 after this timepoint (starting at 27 d old), and two additional groups either fed RU486 from age 15 d until death (‘whole life’) or fed control food (control, see methods and Figure 1). This age-dependent treatment regime was chosen based on an initial experiment measuring age-specific mortality in response to whole life mTor knockdown, as well as a lower sample size experiment that showed mTor knockdown in late life reduced mortality (both unpublished).

**Figure 1.**
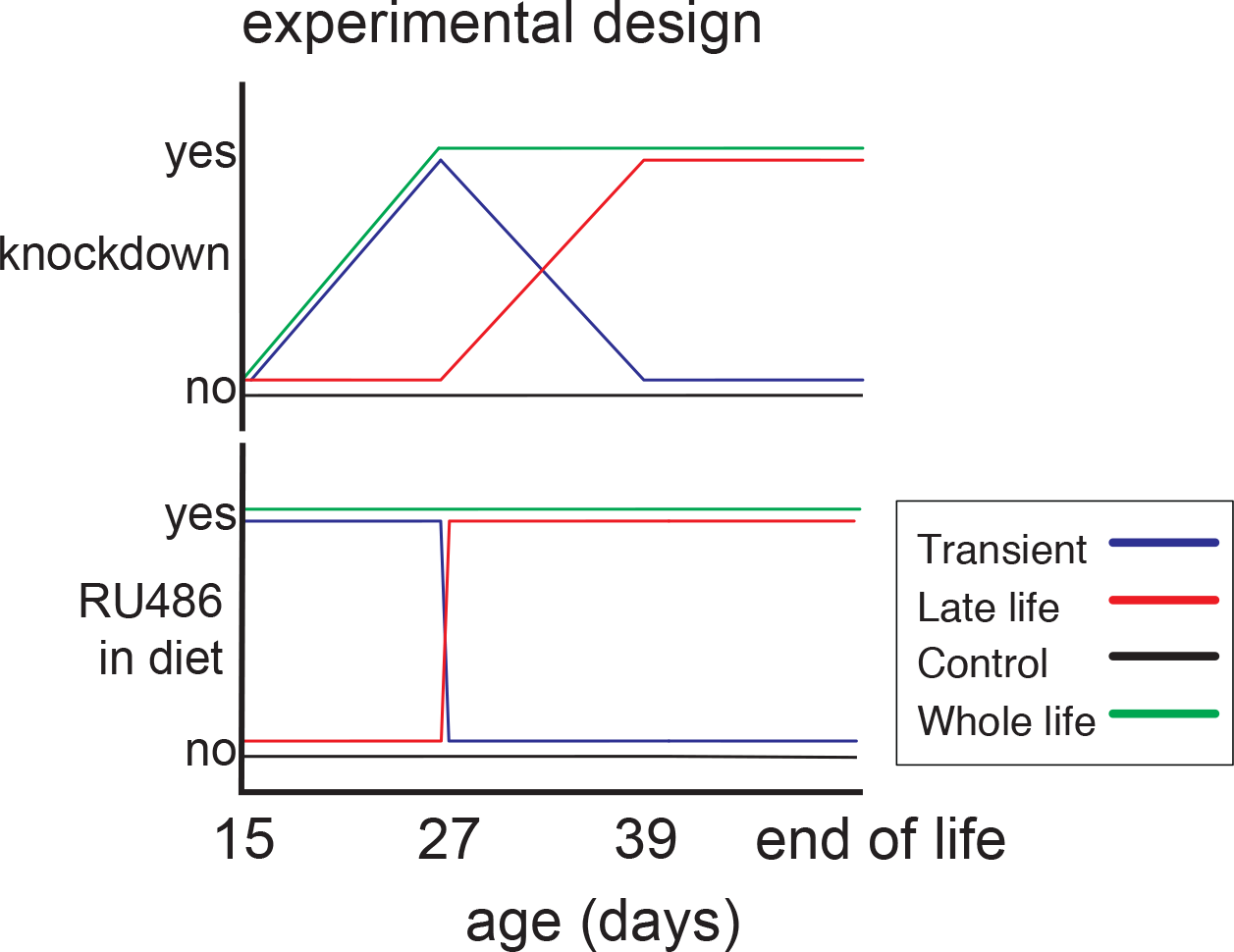
Experimental design. RU486 is fed in an age-dependent manner. Knockdown of mTor is will follow up to maximum induction which is reached after 12 days (Figure 2A). Temporal dynamics of knockdown induction is drawn as linear but could follow other dynamics. Key for interpretation of both mortality and RNAseq data, below, is that maximum response in mortality and associated physiological changes is expected at age 27 and 39.

Control experiments confirmed there were no age-dependent changes in the inducibility of the GeneSwitch system (e.g. due to differential feeding or metabolism of RU486) and determined the kinetics of GeneSwitch induction and termination upon RU feeding. With *daughterless* GeneSwitch (da-GS, expressed in the whole fly)^23^ crossed to overexpression constructs of *hid* and *reaper* (inducers of cell death^26^), we drove and measured the corresponding impact on death relative to RU feeding. Irrespective of age of induction, mortality started to rise after about 2-4 days after initial RU feeding, reached maximum induction after 12 days of RU feeding, and returned to control levels after about 12 day once RU feeding was terminated (Figure 2A, P < 0.0001, N = 3,287). These RU induction dynamics were used to interpret the kinetic effects of mTor knockdown on mortality and transcriptional profiles (Figure 1).

**Figure 2.**
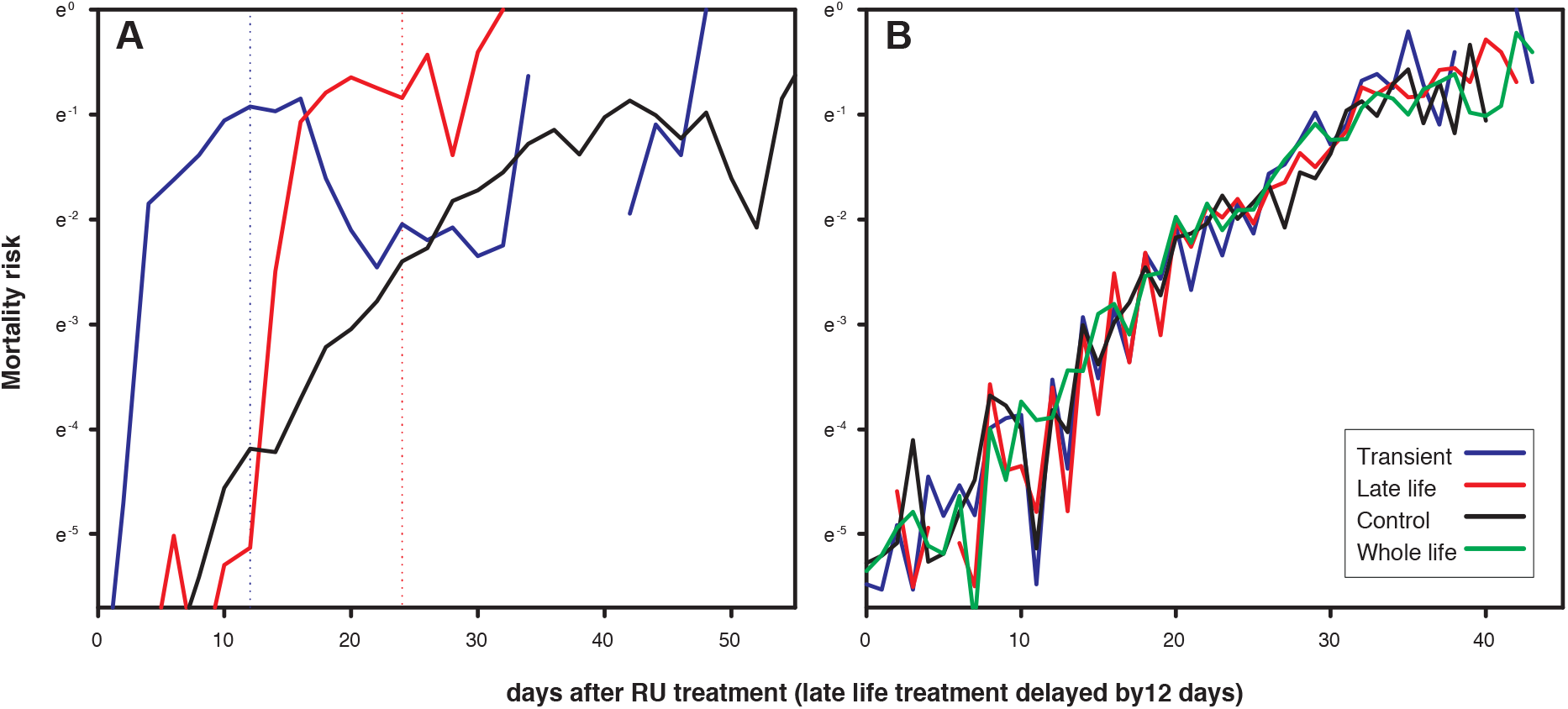
Raw mortality data **A)** *da*-GS (whole fly expression) driving overexpression of *hid* and *rpr* conditionally induced death in the fly, and such induction was reversible in the transient treatment. After the start of RU486 feeding it takes 12 days for full induction of mortality and this drops back to normal levels after removal of RU (see Figure 1 for experimental design). Note control mortality is similar to controls in panel B, suggesting da-GS does not cause any leaky expression, i.e. overexpression of *hid* or *reaper* in the absence of RU486. **B)** Negative control: *da*-GS crossed to a TRiP control line (as dTor knockdown, see below, uses TRiP). In none of the timing regimes did RU486 have any effect on mortality (P > 0.56), as also reported previously in other studies.

### Negative Control

Although no confounding effects of RU486 on mortality have been reported in similar experiments^23,27–30^ we conducted negative controls to exclude any such effects. The da-GS driver line was crossed to a genetic control from the TRiP collection that included the insertion vector without shRNA. Offspring of this cross were studied directly alongside the kinetic control and age-dependent knockdown experiments. When RU486 was applied in the same regime as all induction experiments, no difference in mortality was seen among any treatment (Figure 2B, P > 0.56, N = 4,708).

### Age-dependent effects of mTor knockdown on mortality

Knockdown of mTor at old age instantly decreased mortality (Figure 3A, Hazard rate (exp) = 0.65 ± 0.06, P < 0.0001). Continuous knockdown and transient knockdown of mTor produced similar mortality trajectories where death rate initially increased relative to control but then reduced to a level below that of the control for the remainder of the trial (Figure 3B, Hazard rate (exp) = 0.41—0.43 ± 0.06, P < 0.0001). Because early mortality was induced by mTor knockdown, we evaluated if subsequent lifespan extension could be explained by demographic selection acting on phenotypic heterogeneity for frailty. If the frailest flies were killed by the initial induction of mTor RNAi, mortality measured in the remaining cohort could be reduced by the early loss of this frail subset. To determine if demographic selection accounts for our observed late-life mortality pattern, we generated simulated life tables under an assumption that the excess deaths introduced in early life (N = 329) upon induction of mTor RNAi did not act on frailty variation. We simulated life tables beginning at age 39 that now included the 329 individuals that were lost before age 33 days old (as compared to control) in the observed data by uniformly sampling and reintroduced simulated deaths across this age-interval (ages 39 to 64 days). Note that because mortality increases exponentially, uniform sampling actually biases mortality towards earlier ages and thus simulates the bias induced by phenotypic heterogeneity in frailty. Subsequently we gradually moved and shortened the interval of reintroduction of simulated deaths to earlier ages in subsequent simulations, iteratively increasing demographic selection up to a maximum of all deaths reintroduced at age 39 days. Across all these simulated scenarios, mortality at late ages continued to be significantly reduced in the treatments with early and continuous mTor RNAi (P < 0.0001). Thus, transient knockdown of mTor during early life appears to reduce later mortality through long-lasting physiological effects.

**Figure 3.**
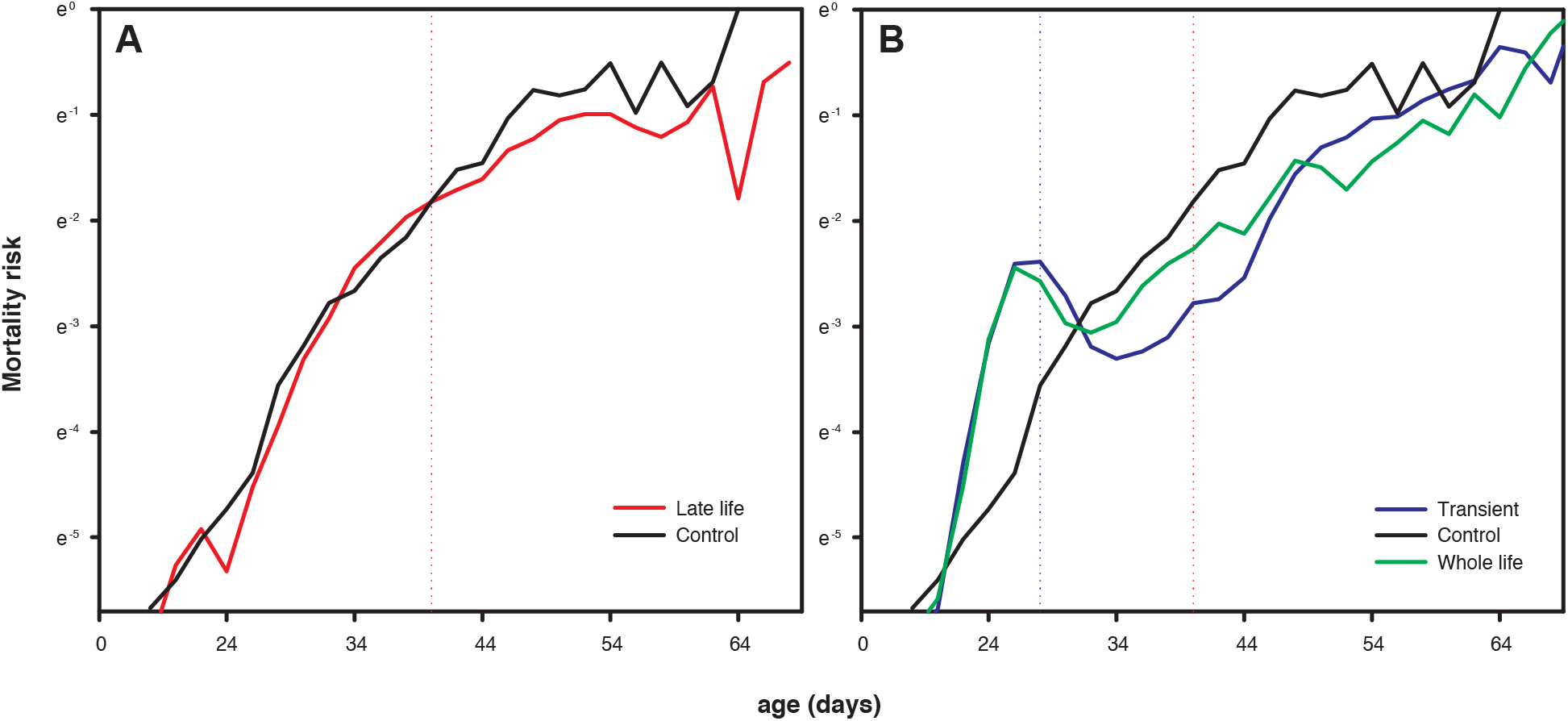
Raw mortality risk plotted on a natural log axis. A) Mortality risk is lowered when mTor is knocked down in late life (P < 0.0001). The red dotted line indicates when da-GeneSwitch is maximally inducing *in vivo* RNAi to knockdown mTor. Data analysed using age-dependent mixed effects cox proportional hazard models correcting for cage effects (see methods^14,31^). B) Transient mTor knockdown (maximum induction of GeneSwitch at blue dotted line) resulted in a modest increase in mortality during early life, but subsequently resulted in a sustained mortality reduction throughout life (P < 0.0001). Similar effects were seen when mTor was knocked down continuously. The red dotted line now indicated when mTor in the transient treatment is back to control levels. These experiments were all ran together at the same time but are split in two panels for graphical purposes.

### Transcriptomes of age-dependent mTor knockdown suggests distinct physiological responses to early and late induction

Conditional knockdown of mTor impacts whole fly mRNA profiles with immediate, reversible and long-lasting changes (Figure 4). Notably, transient mTor reduction induced long-lasting changes in the transcriptome that are distinct from late and whole-life mTor reduction. We used a combined statistical framework (see methods) to distil transcriptional changes that are uniquely associated with the different age-dependent treatment regimes of mTor RNAi (Figure 1). We plotted these effects on the KEGG mTor network to visualize these differences. Knockdown of mTor in late life produced only a limited response across the whole network, whereas transient knockdown in early life induced persistent, substantial differential expressed across the network even though mTor expression was back at control levels (Figure S1).

**Figure 4.**
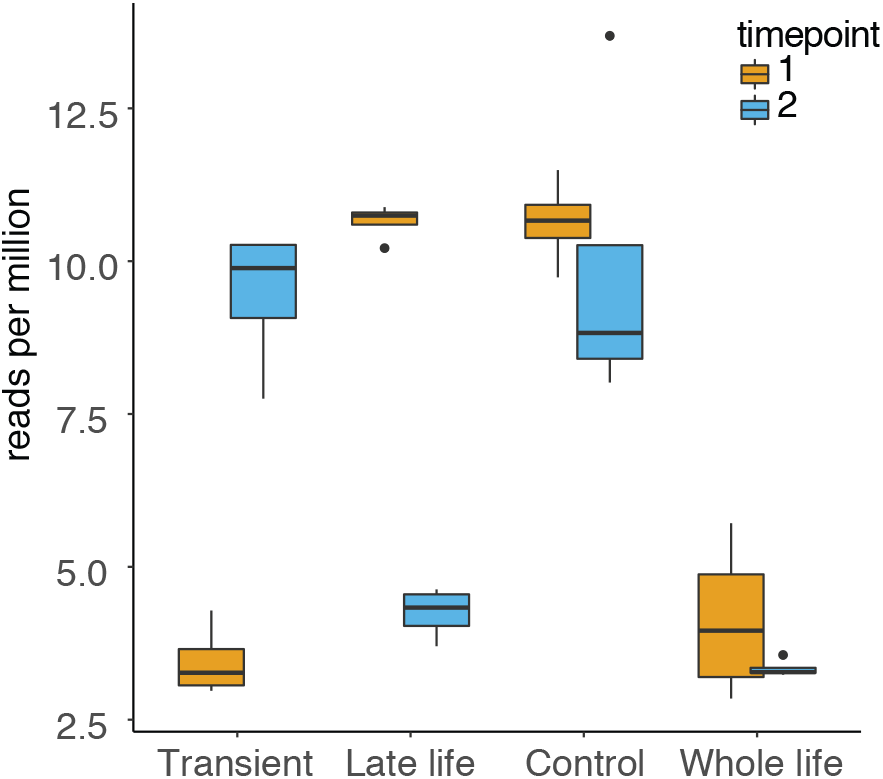
mTor was transiently knocked down, most notably to similar levels in the different treatment categories (F_3,24_= 37.3, P < 0.0001). Blue boxplots indicate the level of mTor expression in RNA from whole flies collected at late life induction (corresponding to red dotted line Figure 3A, see Figure 1). Orange boxplots indicate the same but for early life maximal induction (corresponding to blue line Figure 3B). Transient treatments resulted in immediate experimental changes in transcription in mTor as intended and transient treatment at the respective timepoints where statistically indistinguishable from continuous treatments (P > 0.26).

In the early, transient mTor knockdown cohort, over 6,000 of 10,187 identified transcripts were statistically differentially expressed when compared at old age (model comparison second sampling point at 39 days old, after false-discovery rate Benjamini-Hochberg correction) (Figure 5). Continuous knockdown resulted in over 2,000 differentially expressed genes, whereas late life knockdown resulted in limited differential transcription of around 700 genes. Note, that the comparisons of interest statistically identified here are the additional contribution of the timing of mTor knockdown (early versus late) on top of any effects of whole life knockdown compared to control (for statistical framework see methods). Remarkably, there was only limited overlap in altered transcripts among the treatment categories suggesting the age-dependent effects of mTor knockdown were distinct rather than additive (Figure 5).

**Figure 5.**
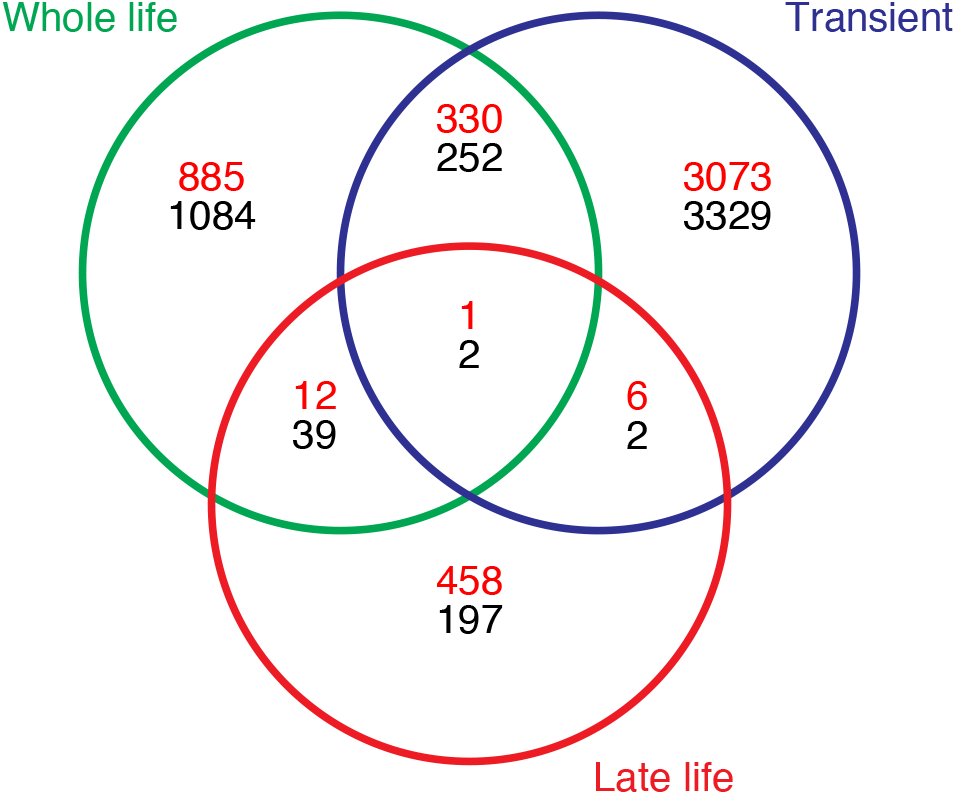
Venn diagram of differentially expressed genes, following false discovery rate correction, compared to the control treatment. Numbers in red are downregulated, black are upregulated, compared to control conditions. Note there is limited overlap between categories suggesting that differences in the transcriptome induced by age-dependent mTor reduction, compared to control, are not additive between treatments, but distinct.

### Age-dependent reduction of mTor signalling implicates different physiology

We performed directional KEGG and GO enrichment analyses^32^ to provide a generalised view of physiological changes between the transcriptional responses across the three distinct timings of mTor knockdown compared to control (Tables S1-S6). Divergent processes were associated with each of the three timing regimes of mTor reduction.

Whole life mTor reduction was associated with upregulated sugar-based metabolism and amino-acid metabolism (Table S1). In contrast to previous studies where mTor suppression upregulates protein processing and the proteasome^33,34^, reduced mTor here downregulated these protein degradation pathways. Variation in temporal dynamics may explain these differences: when the cell is starved it will activate autophagy and protein degradation related mechanisms, but once excess proteins are recycled, the resultant effect is a downregulation of the protein recycling machinery and a shift in total metabolism. The conclusion of the effects of reduced mTor signalling on protein processing and the proteasome might therefore depend on the timing after which this is measured and may depend on the system in which this is studied: cell culture versus whole organism, but this interpretation will require future testing. In line with the interpretation that metabolism is downregulated when mTor is reduced for a prolonged time is that DNA replication and RNA transport are downregulated (Table S1).

In contrast to continuous mTor knockdown, transient depletion of mTor in early adult life upregulated cellular activity at terms representing DNA replication, RNA transport and basal transcription factors, and for ubiquitin mediated proteolysis at old age. Lysosome-associated terms are downregulated as are as elements for metabolism, mainly related to fats (Table S2). We find terms for the spliceosome are widely upregulated, which may explain why transient mTor knockdown has systemic long-term effects on mortality rate and gene expression (Table S2). Notably, work with C. elegans recently suggests that splicing regulation is a potential key mediator of ageing during dietary restriction and by control of mTor^35–38^. We detected limited KEGG enrichment with genes uniquely differentially expressed in the late life mTor knockdown regime suggesting these effects are similar to knockdown of mTor throughout life. Of note, is the partial downregulation of the spliceosome (15 out of 115 genes in this category, Table S3) compared to upregulation in the transient mTor knockdown treatment (84 out of 115). These interpretations based on KEGG were likewise seen in GO analysis, although at lower resolution (Tables S4–S6).

### Alternative splicing

Noting that the spliceosome was upregulated in the transient mTor group, we analysed the RNAseq data set for alternative splicing using exon-level based reads. In the transient mTor knockdown group, 327 genes were significantly differentially spliced, compared to 17 of continuous mTor knockdown and 4 specific to mTor knockdown in late life. KEGG analysis in the transient mTor knockdown group were enriched for differentially spliced variants associated with endocytosis^39^, the lysosome^40^, Hippo signalling (potentially mediating longevity through autophagy^41^ and FOXO^42^) and mTor signalling itself (Table S7). Thus, potentially the long-lasting mortality benefits from transient mTor reduction are mediated by long-lasting changes in alternative splicing, as predicted from the upregulation of the spliceosome^43^.

## Conclusion

Short-term treatments that extend lifespan will be key to translate findings from the field of ageing biology to actual medical applications. These experiments provide the evidence, together with earlier findings from late-life and transient treatment with rapamycin (inhibitor of mTor) in mice^9,18^, that short-term or timed suppression of mTor signalling can have beneficial effects on life/health-span^4^ – a treatment regime that might be practical for humans. While the magnitude of reduced mortality produced by mTor inhibition are less than those we report for diet restriction in Drosophila^1,13^, they are large relative to those gained in humans when key environmental factors are modulated, such as by cessation of smoking^44^. Furthermore, long-term mortality benefits of early, transient mTor depletion appears to operate through different transcriptional changes compared to how mTor affects older animals, and the early impacts appear to involve alternative splicing. The long-lasting benefits from transient treatment could arise from metabolic or signalling reprogramming or hormesis^45^.

These novel insights will help inform future application and potential side effects of drugs targeting mTor. Future work will benefit from uncovering if tissue-specific effects of mTor signalling underlie these age-dependent dynamics and will also need to experimentally test which physiological mechanisms hypothesised from the transcriptome profiles cause the observed mortality differences. These mechanisms will be complex because our data suggest that reduced mTor throughout life probably affects a composite of two (or more) age-dependent processes.

## Methods

### Fly husbandry and mortality measurement

Flies were grown and kept on our standard rich diet (8% autolysed yeast, 13% table sugar, 6% cornmeal, 1% agar and nipagin 0.225% [w/v], with only growing bottles containing 0.4% [v/v] propanoic acid)^14^. Media for vials was cooked and then spilt to allow preparation of drug or control food from the same batch of fly food, controlling for variation in cooking batch. RU486 (Generon UK, dissolved in absolute ethanol at a stock solution of 10mg/ml) was added to the media, during cooling for dispensing into vials, to give a final concentration of 200μM in the fly food. An equal volume of absolute ethanol was added to the control food.

Flies for experiments were grown in controlled density bottles by using a set amount of 10 virgins mated with males. Offspring of each cross was age standardised by transferring all newly eclosed adult flies each day to a new bottle (Flystuff square bottles) for two days of subsequent mating. After mating, flies were sorted into vials, using light CO_2_ anaesthesia, of 25 females each and transferred the same day to a demography cage to contain 125 flies each (for a detailed description see^14^). The cage design^12^ allows the removal of dead flies and changing of food, every other day, without physically transferring the flies, to allow for as little disturbance to them as possible. A low frequency of accidental deaths (stuck to the food or killed accidentally) and escapees were righthand censored in the analyses.

### Statistical analysis of mortality

All mortality data was analysed using mixed effects cox-proportional hazard models using age-dependent covariates^46^ to test for the age-dependent changes in mortality. These models are conservative as they correct for pseudoreplication cause by cage effects (similar to vial to vial error and critical in reducing type I error). Age-dependent covariates allow a comparison during specific ages, specifically relevant as the experimental design induced age-dependent modulation of mTor (Figure 1)^14^. Coefficients are reported in the text and are in comparison to the control treated flies. These coefficients are shown on the linear scale, with standard errors on the log-scale to maintain symmetric errors. Models also included transfer day in the model to correct for any variation between day or growing conditions (although omission of this correction did not change any of the results). All experiments had a maximum of four transferring days into the cages and ages at RU486 treatment thus maximally differed 1-2 days around the mean age of induction. The x-axis, age, in the raw mortality graphs is corrected for such differences in transfer data as all flies (across all the data presented) were given RU486 on the same calendar date. In the graphs this allows an appreciation of the changes in mortality in response to RU486 timing. All analyses included actual non-corrected ages.

### Transcriptome

RNA was extracted from a lysate (generated by bead milling) of ~4 whole flies per sample (4 flies per treatment per timepoint, total 32 samples) using a Qiagen RNeasy mini kit. Samples were shipped on dry ice to the Oxford Genomics Centre where samples were reverse transcribed and an equal concentration was polyA enriched library prepped and deep-sequenced in full multiplex using Illumina HiSeq4000 with 75bp paired ends. Average mapped reads across samples was 176 million. Reads were mapped to the Drosophila melanogaster genome (Release 6) using annotated (and thus not *de novo* assembled) transcripts using *hisat2*^47^, formatted into read counts using *stringtie*^47^ to be used in *ballgown*^47^ in R, and analysed for differential expression using *edgeR* using the general linear modelling framework in *glmQLFit*. We used a full model design correcting for age-dependent changes of mTor knockdown to distil the effects of the timing regimes specifically at late age on the total transcriptome (y ~ knockdown * timepoint + timing regime at old age). This statistical framework therefore identifies differential transcription specific to the timing regime by which mTor is conditionally knocked down compared to control conditions when mortality benefits where observed (Figure 3). Differential splicing was analysed using exome mapping and the function *diffSpliceDGE* from *edgeR*. KEGG and GO enrichments^32^ were conducted using the *limma*^48^ package in R. For plotting of the KEGG mTor pathway, the pathway was updated manually using the most recent fly literature for plotting purposes only (but not for enrichment analyses).

## Acknowledgements

MJPS is supported by a Sir Henry Wellcome and a Sheffield Vice Chancellor’s Fellowship, the Natural Environment Research Council (M005941 & N013832) and an Academy of Medical Sciences Springboard Award (the Wellcome Trust, the Government Department of Business, Energy and Industrial Strategy (BEIS), the British Heart Foundation and Diabetes UK). MT is supported by American Federation of Aging Research, Grant/Award Number: GR5290420; National Institute on Aging, Grant/Award Number:R37 AG024360, T32 AG 41688. ST was supported by the EU Erasmus exchange program with the VU University of Amsterdam. JT was supported by a summer stipend from the University of Oxford. We thank the Oxford Genomics Centre at the Wellcome Centre for Human Genetics (funded by Wellcome Trust grant reference 203141/Z/16/Z) for the generation and initial processing of sequencing data.

---

**Supplementary Figure S1.**
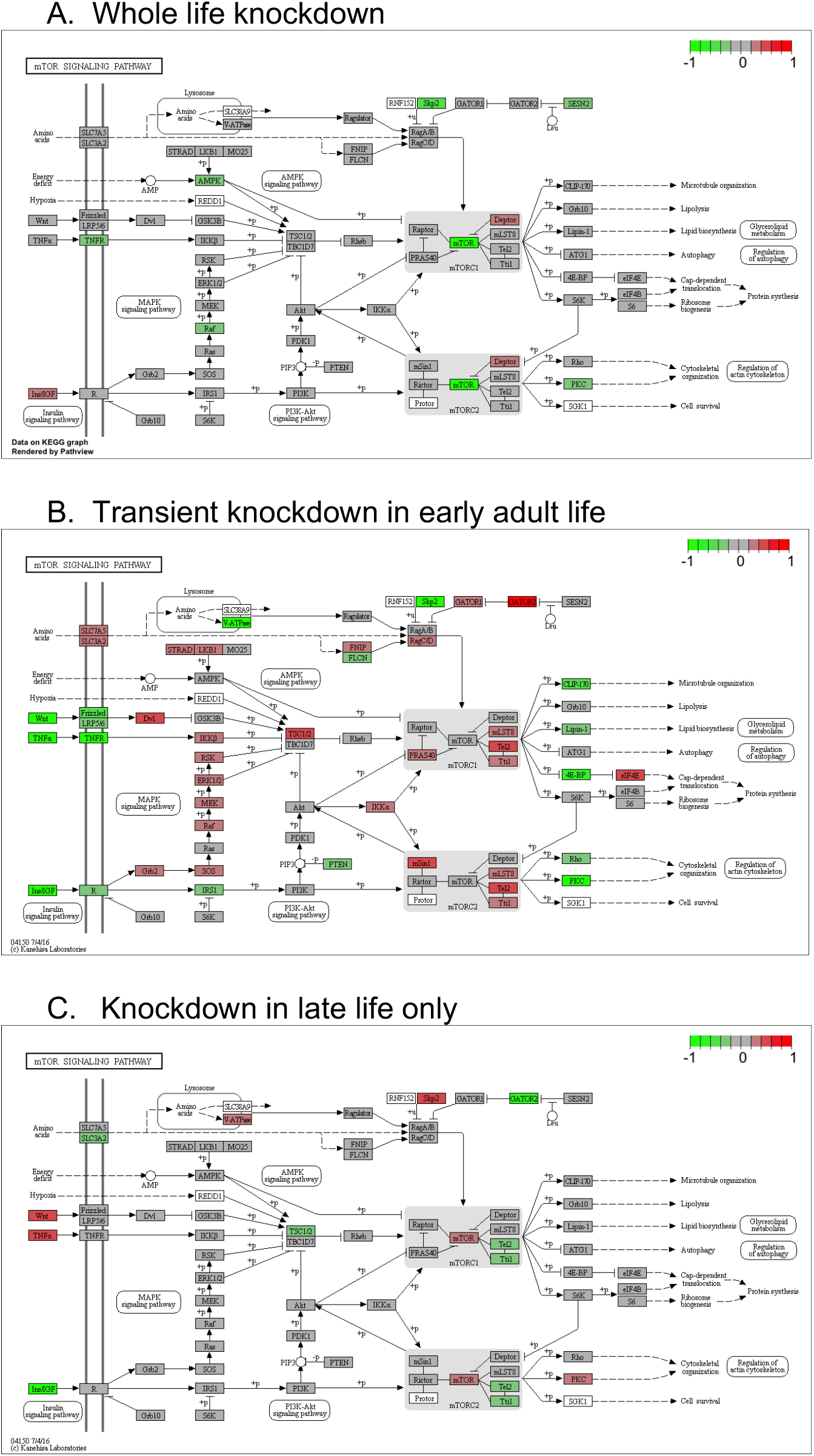
Differential expression of mTor knockdown age-dependent regimes plotted on the wider KEGG mTor network (at late age). Intensity of greener colours indicate reduced transcription relative to control. Intensity of red colours indicate increased transcription relative to control. White boxes indicate genes for which there is no clear paralog in *Drosophila melanogaster*.

**Table S1.** KEGG analysis of transcriptome resulting from mTor knockdown

| Pathway | N | Up | Down | P Up | P Down |
| --- | --- | --- | --- | --- | --- |
| Metabolic pathways | 912 | 172 | 86 | **<0.001** | 0.996 |
| Galactose metabolism | 29 | 13 | 3 | **<0.001** | 0.697 |
| Starch and sucrose metabolism | 28 | 12 | 0 | **<0.001** | 1 |
| Toll and Imd signaling pathway | 63 | 20 | 6 | **<0.001** | 0.788 |
| Carbon metabolism | 91 | 25 | 11 | **<0.001** | 0.546 |
| Glycolysis / Gluconeogenesis | 40 | 14 | 7 | **<0.001** | 0.201 |
| Pentose and glucuronate interconversions | 42 | 14 | 4 | **0.001** | 0.763 |
| Glycine, serine and threonine metabolism | 25 | 10 | 1 | **0.001** | 0.96 |
| Pentose phosphate pathway | 19 | 8 | 2 | **0.002** | 0.687 |
| Drug metabolism - other enzymes | 84 | 21 | 9 | **0.003** | 0.698 |
| Proteasome | 38 | 0 | 33 | 1 | **<0.001** |
| Protein processing in endoplasmic reticulum | 115 | 6 | 32 | 0.999 | **<0.001** |
| DNA replication | 34 | 0 | 14 | 1 | **<0.001** |
| mRNA surveillance pathway | 60 | 4 | 15 | 0.971 | **0.004** |
| Ubiquitin mediated proteolysis | 84 | 4 | 19 | 0.998 | **0.005** |
| Synthesis and degradation of ketone bodies | 7 | 0 | 4 | 1 | **0.005** |
| Mucin type O-glycan biosynthesis | 15 | 1 | 6 | 0.888 | **0.006** |
| Nucleotide excision repair | 37 | 1 | 10 | 0.995 | **0.01** |
| RNA transport | 121 | 7 | 23 | 0.998 | **0.017** |
| Mismatch repair | 20 | 0 | 6 | 1 | **0.027** |

**Table S2.** KEGG analysis of transcriptome resulting from Early adult transient mTor knockdown

| Pathway | N | Up | Down | P Up | P Down |
| --- | --- | --- | --- | --- | --- |
| Spliceosome | 115 | 84 | 8 | **<0.001** | 1 |
| DNA replication | 34 | 33 | 1 | **<0.001** | 1 |
| Nucleotide excision repair | 37 | 35 | 1 | **<0.001** | 1 |
| RNA transport | 121 | 81 | 4 | **<0.001** | 1 |
| Basal transcription factors | 37 | 33 | 1 | **<0.001** | 1 |
| Fanconi anemia pathway | 26 | 24 | 0 | **<0.001** | 1 |
| Mismatch repair | 20 | 19 | 1 | **<0.001** | 1 |
| Homologous recombination | 23 | 21 | 0 | **<0.001** | 1 |
| Ubiquitin mediated proteolysis | 84 | 54 | 4 | **<0.001** | 1 |
| mRNA surveillance pathway | 60 | 41 | 4 | **<0.001** | 1 |
| Neuroactive ligand-receptor interaction | 42 | 0 | 37 | 1 | **<0.001** |
| Biosynthesis of unsaturated fatty acids | 21 | 2 | 17 | 0.999 | **<0.001** |
| Lysosome | 97 | 12 | 53 | 1 | **<0.001** |
| ECM-receptor interaction | 12 | 0 | 10 | 1 | **0.001** |
| Insect hormone biosynthesis | 24 | 1 | 16 | 1 | **0.001** |
| Metabolic pathways | 912 | 227 | 348 | 1 | **0.001** |
| Fatty acid metabolism | 49 | 6 | 27 | 1 | **0.001** |
| Ascorbate and aldarate metabolism | 27 | 3 | 17 | 0.999 | **0.002** |
| Glycerolipid metabolism | 39 | 7 | 22 | 0.995 | **0.003** |
| Caffeine metabolism | 5 | 0 | 5 | 1 | **0.004** |

**Table S3.** KEGG analysis of transcriptome resulting from Late adult mTor knockdown

| Pathway | N | Up | Down | P Up | P Down |
| --- | --- | --- | --- | --- | --- |
| Folate biosynthesis | 34 | 5 | 0 | **0.001** | 1 |
| Thiamine metabolism | 16 | 3 | 0 | **0.006** | 1 |
| ABC transporters | 19 | 3 | 0 | **0.01** | 1 |
| Arginine biosynthesis | 13 | 2 | 0 | **0.037** | 1 |
| Fatty acid elongation | 13 | 2 | 0 | **0.037** | 1 |
| 2-Oxocarboxylic acid metabolism | 14 | 2 | 1 | **0.042** | 0.49 |
| Amino sugar and nucleotide sugar metabolism | 40 | 3 | 2 | 0.068 | 0.567 |
| Biosynthesis of unsaturated fatty acids | 21 | 2 | 0 | 0.087 | 1 |
| Longevity regulating pathway - multiple species | 47 | 3 | 0 | 0.099 | 1 |
| Caffeine metabolism | 5 | 1 | 0 | 0.113 | 1 |
| Ribosome biogenesis in eukaryotes | 72 | 0 | 18 | 1 | **<0.001** |
| Aminoacyl-tRNA biosynthesis | 41 | 0 | 10 | 1 | **<0.001** |
| RNA polymerase | 28 | 0 | 7 | 1 | **<0.001** |
| Spliceosome | 115 | 2 | 15 | 0.76 | **<0.001** |
| Pentose phosphate pathway | 19 | 0 | 4 | 1 | **0.011** |
| RNA transport | 121 | 0 | 12 | 1 | **0.011** |
| Homologous recombination | 23 | 0 | 4 | 1 | **0.021** |
| Glycine, serine and threonine metabolism | 25 | 1 | 4 | 0.451 | **0.028** |
| Non-homologous end-joining | 6 | 0 | 2 | 1 | **0.029** |
| Fanconi anemia pathway | 26 | 0 | 4 | 1 | **0.032** |

**Table S4.**
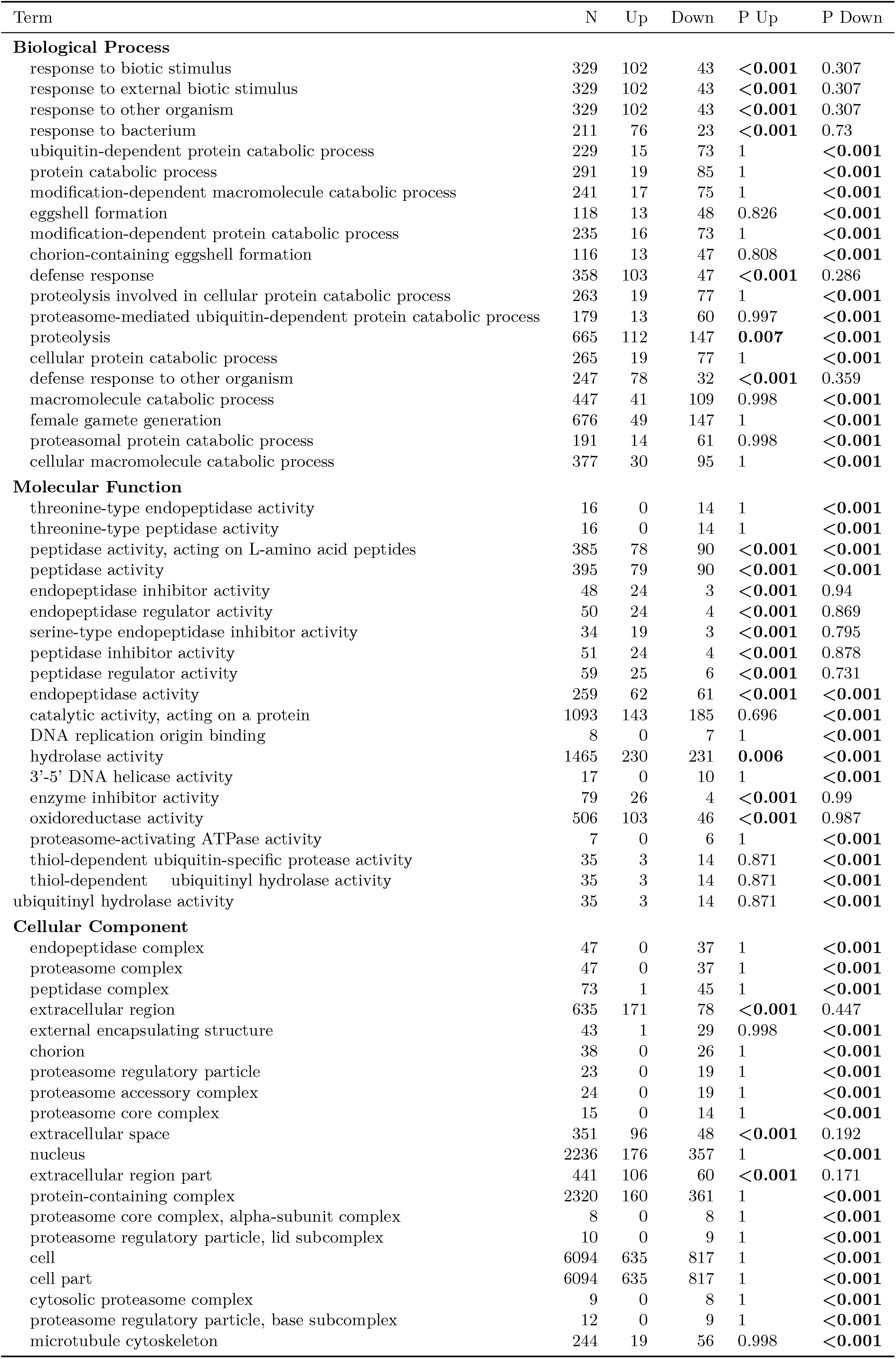
GO analysis of transcriptome resulting from mTor knockdown

| Term | N | Up | Down | P Up | P Down |
| --- | --- | --- | --- | --- | --- |
| **Biological Process** |  |  |  |  |  |
| response to biotic stimulus | 329 | 102 | 43 | <0.001 | 0.307 |
| response to external biotic stimulus | 329 | 102 | 43 | <0.001 | 0.307 |
| response to other organism | 329 | 102 | 43 | <0.001 | 0.307 |
| response to bacterium | 211 | 76 | 23 | <0.001 | 0.73 |
| ubiquitin-dependent protein catabolic process | 229 | 15 | 73 | 1 | <0.001 |
| protein catabolic process | 291 | 19 | 85 | 1 | <0.001 |
| modification-dependent macromolecule catabolic process | 241 | 17 | 75 | 1 | <0.001 |
| eggshell formation | 118 | 13 | 48 | 0.826 | <0.001 |
| modification-dependent protein catabolic process | 235 | 16 | 73 | 1 | <0.001 |
| chorion-containing eggshell formation | 116 | 13 | 47 | 0.808 | <0.001 |
| defense response | 358 | 103 | 47 | <0.001 | 0.286 |
| proteolysis involved in cellular protein catabolic process | 263 | 19 | 77 | 1 | <0.001 |
| proteasome-mediated ubiquitin-dependent protein catabolic process | 179 | 13 | 60 | 0.997 | <0.001 |
| proteolysis | 665 | 112 | 147 | 0.007 | <0.001 |
| cellular protein catabolic process | 265 | 19 | 77 | 1 | <0.001 |
| defense response to other organism | 247 | 78 | 32 | <0.001 | 0.359 |
| macromolecule catabolic process | 447 | 41 | 109 | 0.998 | <0.001 |
| female gamete generation | 676 | 49 | 147 | 1 | <0.001 |
| proteasomal protein catabolic process | 191 | 14 | 61 | 0.998 | <0.001 |
| cellular macromolecule catabolic process | 377 | 30 | 95 | 1 | <0.001 |
| **Molecular Function** |  |  |  |  |  |
| threonine-type endopeptidase activity | 16 | 0 | 14 | 1 | <0.001 |
| threonine-type peptidase activity | 16 | 0 | 14 | 1 | <0.001 |
| peptidase activity, acting on L-amino acid peptides | 385 | 78 | 90 | <0.001 | <0.001 |
| peptidase activity | 395 | 79 | 90 | <0.001 | <0.001 |
| endopeptidase inhibitor activity | 48 | 24 | 3 | <0.001 | 0.94 |
| endopeptidase regulator activity | 50 | 24 | 4 | <0.001 | 0.869 |
| serine-type endopeptidase inhibitor activity | 34 | 19 | 3 | <0.001 | 0.795 |
| peptidase inhibitor activity | 51 | 24 | 4 | <0.001 | 0.878 |
| peptidase regulator activity | 59 | 25 | 6 | <0.001 | 0.731 |
| endopeptidase activity | 259 | 62 | 61 | <0.001 | <0.001 |
| catalytic activity, acting on a protein | 1093 | 143 | 185 | 0.696 | <0.001 |
| DNA replication origin binding | 8 | 0 | 7 | 1 | <0.001 |
| hydrolase activity | 1465 | 230 | 231 | 0.006 | <0.001 |
| 3'-5' DNA helicase activity | 17 | 0 | 10 | 1 | <0.001 |
| enzyme inhibitor activity | 79 | 26 | 4 | <0.001 | 0.99 |
| oxidoreductase activity | 506 | 103 | 46 | <0.001 | 0.987 |
| proteasome-activating ATPase activity | 7 | 0 | 6 | 1 | <0.001 |
| thiol-dependent ubiquitin-specific protease activity | 35 | 3 | 14 | 0.871 | <0.001 |
| thiol-dependent ubiquitinyl hydrolase activity | 35 | 3 | 14 | 0.871 | <0.001 |
| ubiquitinyl hydrolase activity | 35 | 3 | 14 | 0.871 | <0.001 |
| **Cellular Component** |  |  |  |  |  |
| endopeptidase complex | 47 | 0 | 37 | 1 | <0.001 |
| proteasome complex | 47 | 0 | 37 | 1 | <0.001 |
| peptidase complex | 73 | 1 | 45 | 1 | <0.001 |
| extracellular region | 635 | 171 | 78 | <0.001 | 0.447 |
| external encapsulating structure | 43 | 1 | 29 | 0.998 | <0.001 |
| chorion | 38 | 0 | 26 | 1 | <0.001 |
| proteasome regulatory particle | 23 | 0 | 19 | 1 | <0.001 |
| proteasome accessory complex | 24 | 0 | 19 | 1 | <0.001 |
| proteasome core complex | 15 | 0 | 14 | 1 | <0.001 |
| extracellular space | 351 | 96 | 48 | <0.001 | 0.192 |
| nucleus | 2236 | 176 | 357 | 1 | <0.001 |
| extracellular region part | 441 | 106 | 60 | <0.001 | 0.171 |
| protein-containing complex | 2320 | 160 | 361 | 1 | <0.001 |
| proteasome core complex, alpha-subunit complex | 8 | 0 | 8 | 1 | <0.001 |
| proteasome regulatory particle, lid subcomplex | 10 | 0 | 9 | 1 | <0.001 |
| cell | 6094 | 635 | 817 | 1 | <0.001 |
| cell part | 6094 | 635 | 817 | 1 | <0.001 |
| cytosolic proteasome complex | 9 | 0 | 8 | 1 | <0.001 |
| proteasome regulatory particle, base subcomplex | 12 | 0 | 9 | 1 | <0.001 |
| microtubule cytoskeleton | 244 | 19 | 56 | 0.998 | <0.001 |

**Table S5.** GO analysis of transcriptome resulting from Early adult transient mTor knockdown

| Term | N | Up | Down | P Up | P Down |
| --- | --- | --- | --- | --- | --- |
| **Biological Process** |  |  |  |  |  |
| nucleic acid metabolic process | 1858 | 1107 | 259 | <0.001 | 1 |
| cellular macromolecule metabolic process | 2860 | 1519 | 499 | <0.001 | 1 |
| macromolecule metabolic process | 3655 | 1815 | 770 | <0.001 | 1 |
| nucleobase-containing compound metabolic process | 2131 | 1161 | 359 | <0.001 | 1 |
| RNA metabolic process | 1656 | 952 | 252 | <0.001 | 1 |
| heterocycle metabolic process | 2202 | 1184 | 385 | <0.001 | 1 |
| cellular aromatic compound metabolic process | 2259 | 1196 | 414 | <0.001 | 1 |
| cellular metabolic process | 4287 | 1973 | 964 | <0.001 | 1 |
| organic cyclic compound metabolic process | 2309 | 1210 | 432 | <0.001 | 1 |
| gene expression | 2035 | 1096 | 308 | <0.001 | 1 |
| cellular process | 6226 | 2636 | 1688 | <0.001 | 1 |
| cellular nitrogen compound metabolic process | 2586 | 1317 | 450 | <0.001 | 1 |
| nitrogen compound metabolic process | 4076 | 1885 | 957 | <0.001 | 1 |
| primary metabolic process | 4262 | 1926 | 1012 | <0.001 | 1 |
| chromosome organization | 567 | 401 | 32 | <0.001 | 1 |
| regulation of macromolecule metabolic process | 1685 | 907 | 346 | <0.001 | 1 |
| regulation of metabolic process | 1814 | 954 | 381 | <0.001 | 1 |
| regulation of nitrogen compound metabolic process | 1591 | 859 | 335 | <0.001 | 1 |
| cell cycle | 720 | 465 | 68 | <0.001 | 1 |
| regulation of primary metabolic process | 1607 | 864 | 338 | <0.001 | 1 |
| **Molecular Function** |  |  |  |  |  |
| nucleic acid binding | 1464 | 812 | 222 | <0.001 | 1 |
| binding | 4407 | 1894 | 1219 | <0.001 | 1 |
| heterocyclic compound binding | 2319 | 1100 | 521 | <0.001 | 1 |
| organic cyclic compound binding | 2335 | 1101 | 533 | <0.001 | 1 |
| DNA binding | 717 | 421 | 152 | <0.001 | 1 |
| transmembrane transporter activity | 530 | 63 | 313 | 1 | <0.001 |
| inorganic molecular entity transmembrane transporter activity | 378 | 35 | 237 | 1 | <0.001 |
| protein binding | 2023 | 930 | 535 | <0.001 | 1 |
| molecular transducer activity | 208 | 18 | 147 | 1 | <0.001 |
| signaling receptor activity | 208 | 18 | 147 | 1 | <0.001 |
| ion transmembrane transporter activity | 393 | 42 | 235 | 1 | <0.001 |
| transporter activity | 623 | 109 | 333 | 1 | <0.001 |
| transcription factor activity, protein binding | 180 | 129 | 19 | <0.001 | 1 |
| transmembrane signaling receptor activity | 169 | 14 | 120 | 1 | <0.001 |
| chromatin binding | 186 | 131 | 17 | <0.001 | 1 |
| transcription factor activity, transcription factor binding | 166 | 116 | 19 | <0.001 | 1 |
| channel activity | 147 | 12 | 103 | 1 | <0.001 |
| passive transmembrane transporter activity | 147 | 12 | 103 | 1 | <0.001 |
| G-protein coupled receptor activity | 82 | 2 | 67 | 1 | <0.001 |
| cation transmembrane transporter activity | 267 | 28 | 160 | 1 | <0.001 |
| **Cellular Component** |  |  |  |  |  |
| nucleus | 2236 | 1370 | 291 | <0.001 | 1 |
| intracellular part | 5305 | 2521 | 1090 | <0.001 | 1 |
| intracellular | 5350 | 2534 | 1109 | <0.001 | 1 |
| intracellular membrane-bounded organelle | 3636 | 1870 | 632 | <0.001 | 1 |
| intracellular organelle | 4273 | 2102 | 811 | <0.001 | 1 |
| organelle | 4328 | 2115 | 837 | <0.001 | 1 |
| membrane-bounded organelle | 3844 | 1933 | 709 | <0.001 | 1 |
| nuclear part | 1154 | 781 | 58 | <0.001 | 1 |
| intracellular organelle part | 2481 | 1317 | 319 | <0.001 | 1 |
| cell | 6094 | 2628 | 1562 | <0.001 | 1 |
| cell part | 6094 | 2628 | 1562 | <0.001 | 1 |
| organelle part | 2542 | 1327 | 352 | <0.001 | 1 |
| protein-containing complex | 2320 | 1232 | 333 | <0.001 | 1 |
| intracellular organelle lumen | 1012 | 631 | 57 | <0.001 | 1 |
| membrane-enclosed lumen | 1012 | 631 | 57 | <0.001 | 1 |
| organelle lumen | 1012 | 631 | 57 | <0.001 | 1 |
| nuclear lumen | 827 | 539 | 43 | <0.001 | 1 |
| chromosome | 521 | 370 | 31 | <0.001 | 1 |
| intrinsic component of plasma membrane | 446 | 42 | 322 | 1 | <0.001 |
| integral component of plasma membrane | 433 | 39 | 315 | 1 | <0.001 |

**Table S6.** GO analysis of transcriptome resulting from Late adult mTor knockdown

| Term | N | Up | Down | P Up | P Down |
| --- | --- | --- | --- | --- | --- |
| **Biological Process** |  |  |  |  |  |
| ncRNA metabolic process | 321 | 0 | 67 | 1 | <0.001 |
| gene expression | 2035 | 9 | 192 | 1 | <0.001 |
| RNA metabolic process | 1656 | 8 | 166 | 1 | <0.001 |
| nucleic acid metabolic process | 1858 | 9 | 178 | 1 | <0.001 |
| ncRNA processing | 245 | 0 | 55 | 1 | <0.001 |
| RNA processing | 569 | 0 | 86 | 1 | <0.001 |
| cellular nitrogen compound metabolic process | 2586 | 18 | 216 | 1 | <0.001 |
| nucleobase-containing compound metabolic process | 2131 | 17 | 187 | 1 | <0.001 |
| heterocycle metabolic process | 2202 | 18 | 189 | 1 | <0.001 |
| organic cyclic compound metabolic process | 2309 | 23 | 193 | 1 | <0.001 |
| cellular aromatic compound metabolic process | 2259 | 21 | 189 | 1 | <0.001 |
| transmembrane transport | 487 | 47 | 10 | <0.001 | 0.999 |
| ribosome biogenesis | 212 | 0 | 43 | 1 | <0.001 |
| rRNA metabolic process | 151 | 0 | 36 | 1 | <0.001 |
| ribonucleoprotein complex biogenesis | 293 | 1 | 50 | 0.999 | <0.001 |
| macromolecule metabolic process | 3655 | 45 | 255 | 1 | <0.001 |
| rRNA processing | 141 | 0 | 34 | 1 | <0.001 |
| nitrogen compound metabolic process | 4076 | 60 | 270 | 1 | <0.001 |
| primary metabolic process | 4262 | 60 | 278 | 1 | <0.001 |
| tRNA metabolic process | 131 | 0 | 29 | 1 | <0.001 |
| **Molecular Function** |  |  |  |  |  |
| catalytic activity, acting on RNA | 246 | 0 | 50 | 1 | <0.001 |
| RNA binding | 582 | 0 | 79 | 1 | <0.001 |
| nucleic acid binding | 1464 | 5 | 140 | 1 | <0.001 |
| heterocyclic compound binding | 2319 | 27 | 187 | 1 | <0.001 |
| organic cyclic compound binding | 2335 | 28 | 187 | 1 | <0.001 |
| transmembrane transporter activity | 530 | 46 | 11 | <0.001 | 1 |
| secondary active transmembrane transporter activity | 124 | 23 | 1 | <0.001 | 0.998 |
| transporter activity | 623 | 46 | 19 | <0.001 | 0.986 |
| active transmembrane transporter activity | 229 | 27 | 5 | <0.001 | 0.985 |
| organic anion transmembrane transporter activity | 126 | 20 | 0 | <0.001 | 1 |
| anion transmembrane transporter activity | 157 | 21 | 1 | <0.001 | 0.999 |
| sodium-independent organic anion transmembrane transporter activity | 27 | 10 | 0 | <0.001 | 1 |
| ion transmembrane transporter activity | 393 | 32 | 3 | <0.001 | 1 |
| catalytic activity, acting on a tRNA | 90 | 0 | 20 | 1 | <0.001 |
| helicase activity | 93 | 0 | 20 | 1 | <0.001 |
| chitin binding | 62 | 12 | 1 | <0.001 | 0.95 |
| inorganic molecular entity transmembrane transporter activity | 378 | 29 | 4 | <0.001 | 1 |
| ATP-dependent helicase activity | 71 | 0 | 16 | 1 | <0.001 |
| purine NTP-dependent helicase activity | 71 | 0 | 16 | 1 | <0.001 |
| catalytic activity | 3401 | 85 | 211 | 0.284 | <0.001 |
| **Cellular Component** |  |  |  |  |  |
| nuclear part | 1154 | 2 | 116 | 1 | <0.001 |
| nucleus | 2236 | 10 | 180 | 1 | <0.001 |
| intracellular | 5350 | 71 | 330 | 1 | <0.001 |
| intracellular part | 5305 | 71 | 328 | 1 | <0.001 |
| intracellular organelle lumen | 1012 | 2 | 98 | 1 | <0.001 |
| membrane-enclosed lumen | 1012 | 2 | 98 | 1 | <0.001 |
| organelle lumen | 1012 | 2 | 98 | 1 | <0.001 |
| intracellular membrane-bounded organelle | 3636 | 35 | 244 | 1 | <0.001 |
| nucleolus | 209 | 0 | 36 | 1 | <0.001 |
| nuclear lumen | 827 | 2 | 82 | 1 | <0.001 |
| membrane-bounded organelle | 3844 | 37 | 246 | 1 | <0.001 |
| preribosome | 72 | 0 | 19 | 1 | <0.001 |
| ribonucleoprotein complex | 570 | 0 | 61 | 1 | <0.001 |
| integral component of membrane | 1057 | 55 | 17 | <0.001 | 1 |
| protein-containing complex | 2320 | 17 | 161 | 1 | <0.001 |
| intrinsic component of membrane | 1077 | 55 | 17 | <0.001 | 1 |
| cell | 6094 | 113 | 342 | 1 | <0.001 |
| cell part | 6094 | 113 | 342 | 1 | <0.001 |
| integral component of plasma membrane | 433 | 31 | 3 | <0.001 | 1 |
| aminoacyl-tRNA synthetase multienzyme complex | 10 | 0 | 7 | 1 | <0.001 |

**Table S7.** KEGG analysis of alternative splicing at old age induced by transient mTor knockdown

| Pathway | N | Differentially expressed | P Enrichment |
| --- | --- | --- | --- |
| Endocytosis | 122 | 11 | **0.002** |
| Hippo signaling pathway - fly | 61 | 6 | **0.013** |
| Glycerophospholipid metabolism | 63 | 6 | **0.015** |
| mTOR signaling pathway | 96 | 7 | **0.035** |
| Lysosome | 118 | 8 | **0.036** |

